# Generative pretraining from large-scale transcriptomes: Implications for single-cell deciphering and clinical translation

**DOI:** 10.1101/2022.01.31.478596

**Authors:** Hongru Shen, Xilin Shen, Jiani Hu, Jilei Liu, Chao Zhang, Dan Wu, Mengyao Feng, Meng Yang, Yang Li, Yichen Yang, Wei Wang, Qiang Zhang, Jilong Yang, Kexin Chen, Xiangchun Li

**Affiliations:** Tianjin Cancer Institute, National Clinical Research Center for Cancer, Key Laboratory of Cancer Prevention and Therapy, Tianjin Medical University Cancer Institute and Hospital, Tianjin Medical University, Tianjin, China; Department of Bone and Soft Tissue Tumor, Tianjin Medical University Cancer Institute and Hospital, Tianjin Medical University, Tianjin, China; Department of Epidemiology and Biostatistics, Key Laboratory of Molecular Cancer Epidemiology of Tianjin, National Clinical Research Center for Cancer, Key Laboratory of Cancer Prevention and Therapy, Tianjin Medical University Cancer Institute and Hospital, Tianjin Medical University Cancer Institute and Hospital, Tianjin Medical University, Tianjin, China; Department of Maxillofacial and Otorhinolaryngology Oncology, Tianjin Medical University Cancer Institute and Hospital, Tianjin Medical University, Tianjin, China

**Keywords:** single-cell, transcriptome, generative pretraining, deep learning

## Abstract

Exponential accumulation of single-cell transcriptomes poses great challenge for efficient assimilation. Here, we present an approach entitled *tGPT* towards integration of 22.3 million single-cell transcriptomes by modeling gene expression rankings as generative pretraining task. *tGPT* is conceptually simple in that it autoregressively models the ranking of a gene in the context of its preceding neighbors. We demonstrated the high performance of *tGPT* on a range of fundamental single-cell analysis tasks and novel applications on bulk tissues. The single-cell clusters and cell lineage trajectories derived from *tGPT* are highly aligned with known cell labels and states. The feature patterns of tumor bulk tissues learned by *tGPT* are associated with a wide range of genomic alteration events, prognosis and treatment outcome of immunotherapy. *tGPT* represents a new analytical paradigm for integrating and deciphering massive amount of transcriptome data and it will facilitate the interpretation and clinical translation of single-cell transcriptomes.

## Introduction

Rapid advancement in single-cell RNA sequencing leads to dramatical drop in sequencing cost and allows for millions of single-cell transcriptomes to be digitized in a single experiment simultaneously. The whole human body is estimated to have 30 trillion cells. Single-cell transcriptome sequencing provided an unprecedented resolution to distinguish different cell type clusters, depict hierarchical cell arrangement and decipher transitional cell states. To achieve this goal, multiple single-cell atlasing projects have been established internationally, including Human Cell Atlas (HCA)^1^, Single Cell Expression Atlas (SCEA)^2^, COVID-19 Atlas^3^, Tabula Muris Atlas^4^ and Mouse Cell Atlas^5^. The HCA project^1^ aims to digitize all cells and create a reference map of the human body through community-driven initiative that researchers all around the world can contribute. SCEA^2^ compiles and annotates publicly available single-cell transcriptomes across multiple species and different studies. The COVID-19 Atlas^3^ aims at elucidating molecular mechanism and therapeutic target of COVID-19 by generating single-cell atlas of SARS-CoV-2 infection in COVID-19 patients. The Tabula Muris^4^ and MCA^5^ atlases constitute the single-cell reference maps of mouse with millions of cells obtained from different organs. These atlasing projects pose tremendous challenge in the integration of diverse transcriptomes from different projects. However, single-cell transcriptomes are generated by different platforms and experimental protocols. They are sparse, noise and prone to batch-effect^6,7^. Therefore, an analytical method to efficiently integrate ten millions of cells are urgently needed.

Over the past few years, deep learning approaches have led to seismic changes in image recognition and natural language understanding. The success of deep learning could largely attribute to the availability of big data, advancement in computational infrastructure, expressivity and scalability of the computational model. The deep learning model could adeptly handle super large-scale high dimensional data and assimilate real-world information. Due to the exponential accumulation of millions of cell transcriptomes, elucidation of the reference map of single-cell transcriptomes with deep learning becomes an attractive application. Deep learning methods such as *scVI*^8^, *SAUCIE*^9^ and *INSCT*^10^ have been developed for the analysis of single-cell transcriptomes.

The progress of artificial intelligence is undergoing a paradigm shift in computer vision and natural language processing. Deep neural networks based on transformer are becoming the de facto approach in wide variety of scenarios such as vision, language and reasoning^11^. Transformer-based models pretrained on broad data at scale continues to achieve state-of-the-art progress in image classification^12,13^ and language understanding^14-16^. The success of these pretrained models can be attributed to their high expressivity and scalability enabled by transformer to assimilate feature representation from massive amount of unlabeled data. However, the investigation of single-cell transcriptome pretraining at scale has not been well studied.

In this study, we present a deep learning approach entitled *tGPT* towards integration of unlimited number of cells. *tGPT* is built upon transformer that has been widely used in natural language understanding and image recognition. The transformer is an essential component and key success of foundation models because of its high expressivity and scalability^11^. *tGPT* takes as input the expression rankings of top-expressing genes rather than the actual expression levels. *tGPT* is conceptually simple and empirically efficient. It models the occurrence of a gene in the context of its preceding neighbors’ rankings. We developed *tGPT* with 22.3 million cells and systematically evaluated *tGPT* on several heterogeneous datasets for sensitivity to batch-effect, delineation of clustering performance and inference of developmental lineages. We applied *tGPT* to bulk cancer tissue sequencing samples and found that features obtained from *tGPT* are significantly associated with diverse genomic alteration events, patients’ prognosis and treatment outcome of immunotherapy. *tGPT* represents a new analytical paradigm to integrate and decipher large-scale single-cell transcriptomes. It will facilitate the integration and clinical translation of large volume of single-cell transcriptome data.

## Results

### An overview of *tGPT* and its downstream applications

The analytical framework of *tGPT* (**Figure 1**) consists of three components: development of *tGPT*, applications of *tGPT* for single-cell clustering and inference of developmental lineage and interrogation of feature representation of bulk tissues in relation to genomic alterations, prognosis and treatment response of immunotherapy.

**Figure 1.**
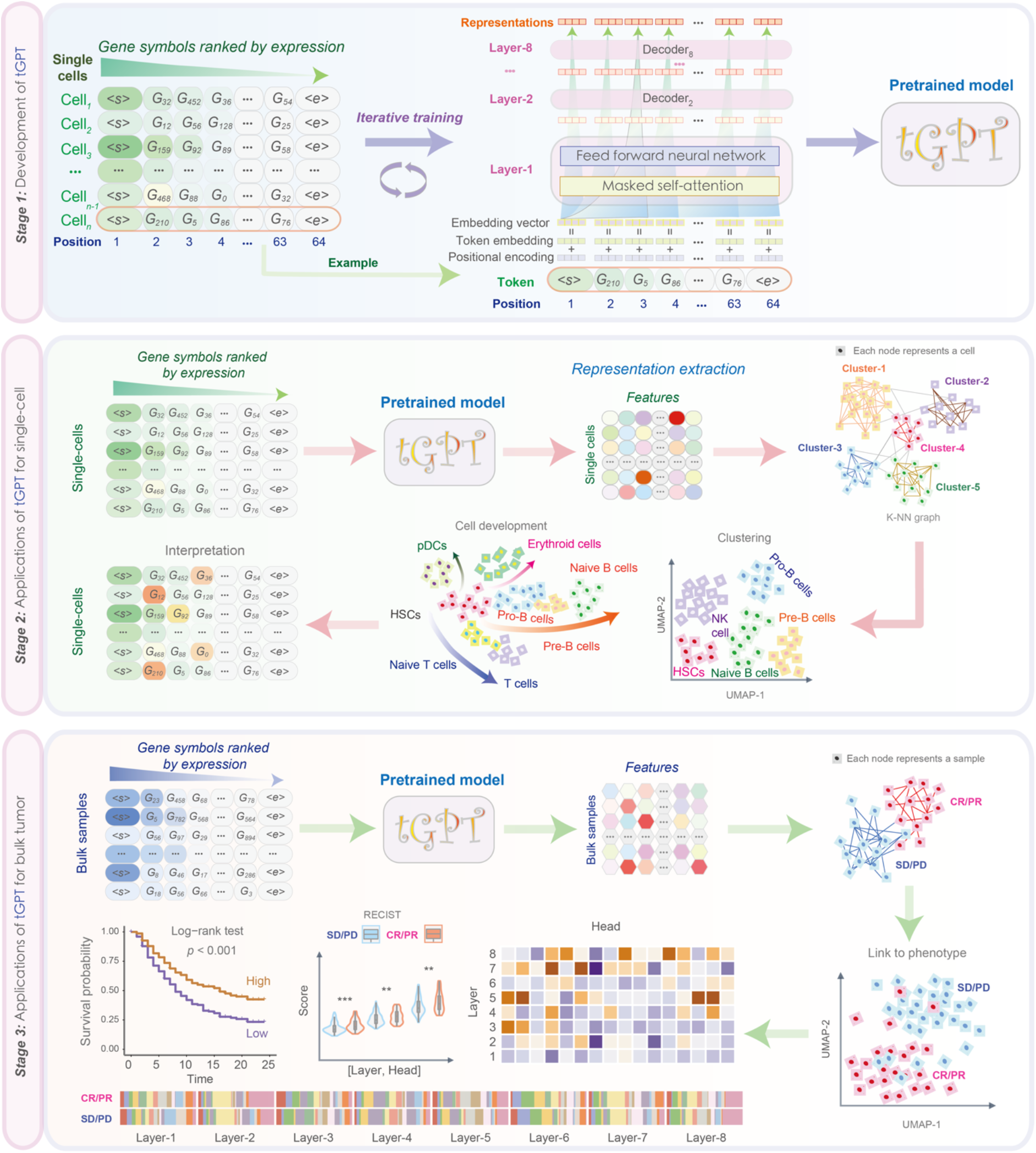
A flowchart illustrating the framework of *tGPT* and its downstream applications. It consists of three components: development of *tGPT*, applications of *tGPT* for single-cell and bulk tissue transcriptomes.

*tGPT* is formatted as an autoregressive language model in that the output from the previous step is used as input to the next step. The input to *tGPT* is a sequence of gene symbols that are ranked by their expression levels. The purpose is to predict the index of the next gene in the dictionary in the context of all previous genes. The dictionary consists of 20706 protein-coding genes. *tGPT* is trained as an unsupervised generative pretraining task^16^. Specifically, for a given cell, let 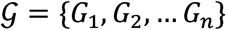 denote the gene symbols that are sorted in a descending order according to their expression levels. We use the standard language modeling objective 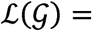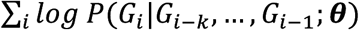 to maximize the likelihood. Here, *k* is the width of context window and **θ** are the parameters of *tGPT* that is used to model the conditional probability. The neural network consists of 8 transformer decoder blocks^17^ with 1024 hidden units and 16 attention heads.

### Quantitative evaluations of *tGPT* on clustering

We systematically evaluated the clustering performance of *tGPT* on four heterogeneous single-cell datasets of different sizes (50-586k cells) from different species and two bulk tissue sequencing datasets. These four single-cell datasets include Human Cell Atlas Census of Immune Cells^18^ (*HCA*, n = 282,558), Human Cell Landscape^19^ (*HCL*, n = 586,135), *Tabula Muris*^4^ (n = 54,862) and *Macaque Retina*^20^ (n = 124,965) dataset (See methods for description). The two bulk tissue datasets are *Genotype-Tissue Expression*^21^ (*GTEx*, n = 11,688) derived from 30 organs and *The Cancer Genome Atlas*^22^ (*TCGA*, n = 9,318) consisted of 33 cancer types.

The clustering performance of *tGPT* is robust with respect to the numbers of top-expression genes being used. We found that the performance of *tGPT* pretrained on the ranking of top 62 and 126 genes were comparable across these six datasets (**Supplementary Figure 1**). In addition, we observed that clustering performance on features extracted from different transformer layers [Layer-1, …, Layer-8] are comparable and better than features extracted from the embedding layer across all these six datasets (**Supplementary Figure 1**). We performed grid-search to identify optimal values of two parameters that are most relevant to clustering (see **Methods**) and reported the best performance for each method. The result from grid-search were provided in **Supplementary Figures 2-7**. Quantitatively, *tGPT* achieved an *NMI* ranged from 0.75 on *HCA* to 0.90 on *GTEx*, *ARI* from 0.53 on *HCL* to 0.84 on *Tabula Muris* and *FMI* from 0.55 on *HCL* to 0.85 on *Tabula Muris* (**Figure 2A**). The clustering performance achieved by *tGPT* are comparable to the other methods such as Scanpy^23^, Pegasus^24^ and scVI^8^ (**Supplementary Figures 8-11)**. Grid-search results of these methods were provided in **Supplementary Figure 12**.

**Figure 2.**
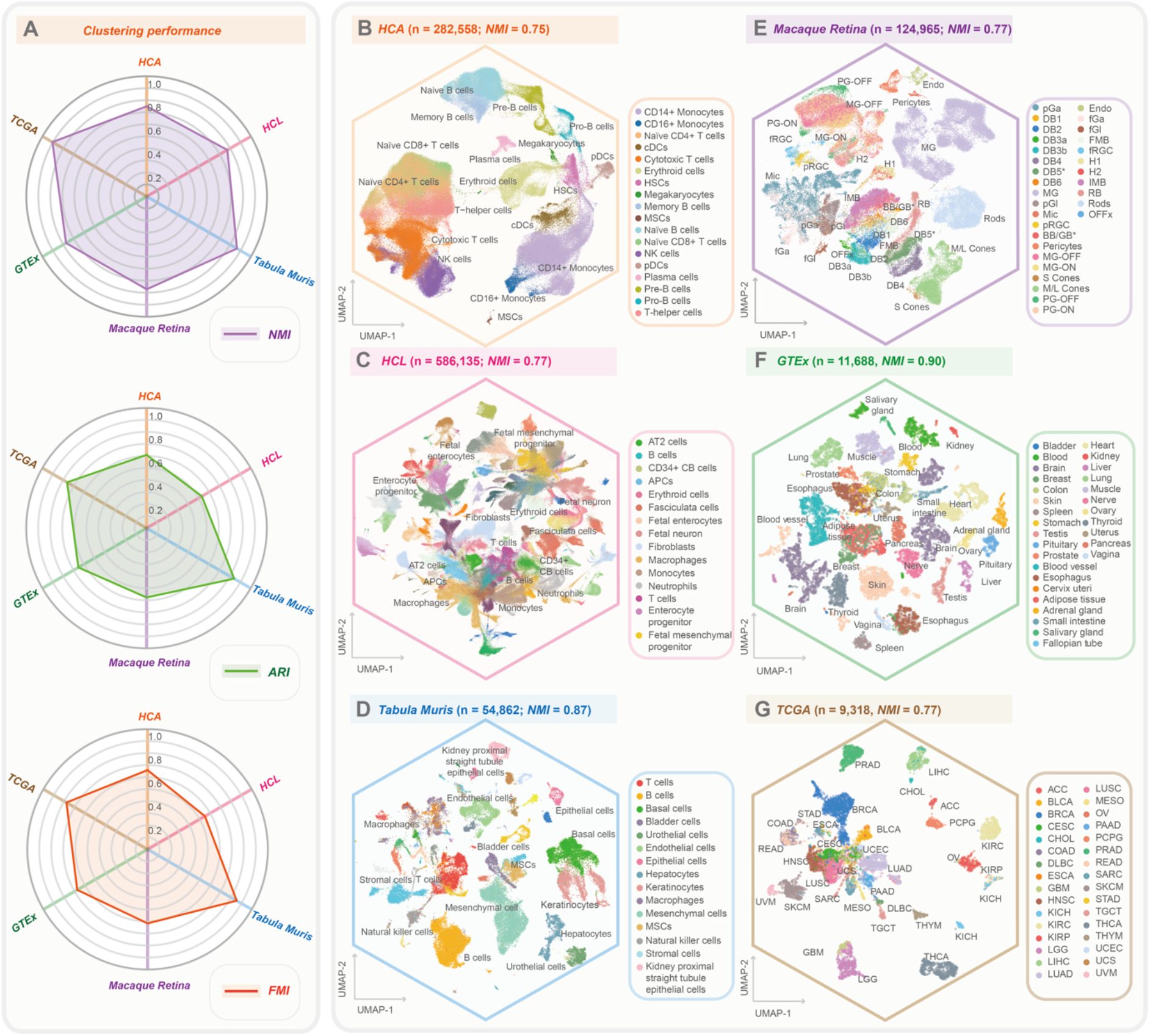
The clustering performance of *tGPT* on four single-cell and two bulk tissue datasets. (A) Radar charts depicting clustering metrics of t*GPT* across these six datasets. UMAP visualization of feature representations learned by *tGPT* on the *HCA* **(B)**, *HCL* **(C)**, *Tabula Muris* **(D)**, *Macque Retina* **(E)**, *GTEx* **(F)** and *TCGA* **(G).** *NMI*, Normalized Mutual information ; *ARI*, Adjusted Rand Index; *FMI*, Fowlkes-Mallows Index.

Across these datasets, *tGPT* was capable of grouping cells with the same or similar types (**Figure 2B-G**). On the *HCA* dataset, *tGPT* was able to identify cells at different developmental phases. For example, it can delineate B cells of different types such as naïve B cells, precursor B (pre-B) cells and progenitor B (pro-B) cells and homologous cells such as conventional DCs (cDCs) and plasmacytoid DCs (pDC), CD14+ and CD16+ monocytes. Less represented cell types such as megakaryocytes (0.32%) and MSCs (0.10%) were also captured by *tGPT* (**Figure 2B**). On the *HCL* dataset, *tGPT* was able to distinguish between immune cells and nonimmune cells as well as different cell types from fetus and adult such as fetal enterocytes and adult enterocytes (**Figure 2C and Supplementary Figure 9**). On the *Tabula Muris* dataset, *tGPT* was also able to delineate 55 distinct cell types originated from 20 mouse organs (**Figure 2D** and **Supplementary Figure 10**). On the *Macaque Retina* dataset, distinctive cell clusters from foveal and peripheral regions of fascicularis retina defined by *tGPT* are well matched with cell types defined in the original literature^20^ (**Figure 2E**). On the *GTEx* dataset, *tGPT* is able to identify different tissues originated from lineage of organs (*NMI* = 0.90), and samples with similar histological structure are close together such as colon, small intestine and stomach (**Figure 2F**). On the *TCGA* dataset, different cancer types are well separated (*NMI* = 0.77). Cancer types with the same tissue of origin tend to clump together in the feature representation spaces captured by *tGPT*. For example, adenocarcinomas and squamous cell carcinomas are closely related in the UMAP plots, respectively.

We observed that *tGPT* is insensitive to batch effect as benchmarked against with the other methods that support batch-correction such as *ComBat*^25^, *MNN*^26^, *Harmony*^27^, *Seurat*^28,29^, *BBKNN*^30^, *Scanorama*^31^, *Pegasus*^24^, *scVI*^8^, *scArches*^32^, *iMAP*^33^ and *DESC*^34^ as measured on the *HCA* dataset. *tGPT* achieved a comparable *kBET* acceptance rate^35^ of 0.87 among the aforementioned batch-correction methods (**Supplementary Figure 13L**). The UMAP plots of these batch-correction methods and their clustering metrics and grid-search results are provided in **Supplementary Figures 13A-K, 14 and 15**, respectively.

### Distinct features learned by *tGPT* are connected to cell types

We observed that the head entropy and importance of different cell types from the *HCA* dataset (See **Methods**) are distinctive from each other. Cells of similar lineages or functions such as T-lineage cells exhibited similar entropy patterns (**Figure 3A**). The head importance is varying considerably for different cell types, however, cells of similar types are alike as compared with the other cell types (**Figure 3B**). For each cell type, we calculated the contribution of each gene on the cell final feature representation (See **Methods**). Cell type specific genes have higher attribution scores (**Figure 3C**). For example, *NKG7*, *FGFBP2*, *PRF1*, *GNLY*, *GZMA* and *GZMB* are highly represented in cytotoxic T cells and NK cells (**Figure 3D**). *PPBP* and *PF4* are also highly represented in megakaryocytes (**Figure 3E**). B-lineage cells have high attribution scores for both *CD79A* and *CD79B.* Attribution scores of *MS4A1* and *MZB1* are relative higher in memory B cells and plasma cells, respectively (**Figure 3F**). The attribution score of *CST3* is higher among CD14+ monocytes, CD16+ monocytes, cDCs and pDCs. In addition, each specific cell types can be defined by specific genes with high attribution scores, for instance plasmacytoid dendritic cells (pDCs, *IRF7*), conventional dendritic cells (cDC, *FCER1A*), CD14+ monocytes (*CD14*) and CD14+ monocytes (*FCGR3A*) (**Figure 3G**).

**Figure 3.**
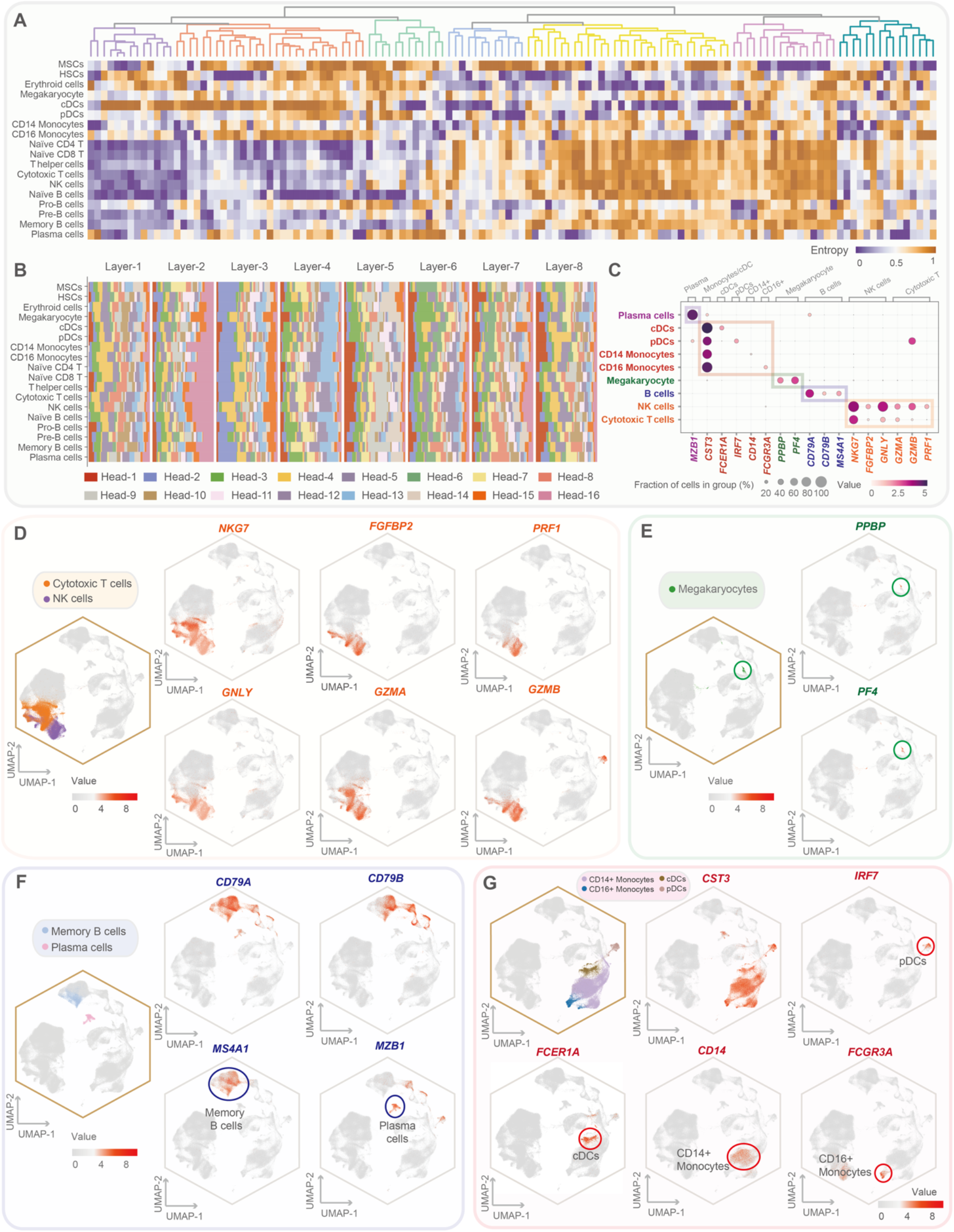
Distinct features of different cell types from the *HCA* dataset learned by *tGPT*. **(A)** Heatmap representation of attention head entropy for different cell types. **(B)** Heatmap representation of attention head importance for different cell types. **(C)** Dot plot illustrating the attribution scores for cell type specific genes. **(D-G)** Scatter plots illustrating the distribution of attribution scores for different cell type specific genes across different cell types.

### Inference of developmental lineage

We used the feature representations learned by *tGPT* to construct cell pseudo-temporal trajectories on *HCA* and *HCL* datasets (See **Methods**). On the *HCA* dataset, the developmental trajectories originated from stem cells and differentiated towards multiple biologically functional cell branches (**Figure 4A**): HSCs to erythroid cells^36^ or DCs and monocytes (**Figure 4B**); naïve T cells to cytotoxic T cells and NK cells^37^ (**Figure 4C**); pro-B cells to pre-B cells, then followed by matured naïve B cells, and finally bifurcated into memory B cells or plasma cells^38^ (**Figure 4D**). In addition, we observed that the cell state signatures are aligned with cell developmental lineages (See **Methods**). For instance, HSCs and pro-B cells are manifested by apparent progenitor signaling (**Figure 4E**). Naïve and mature T cells are featured by distinguishable patterns (**Figure 4F** and **G**).

**Figure 4.**
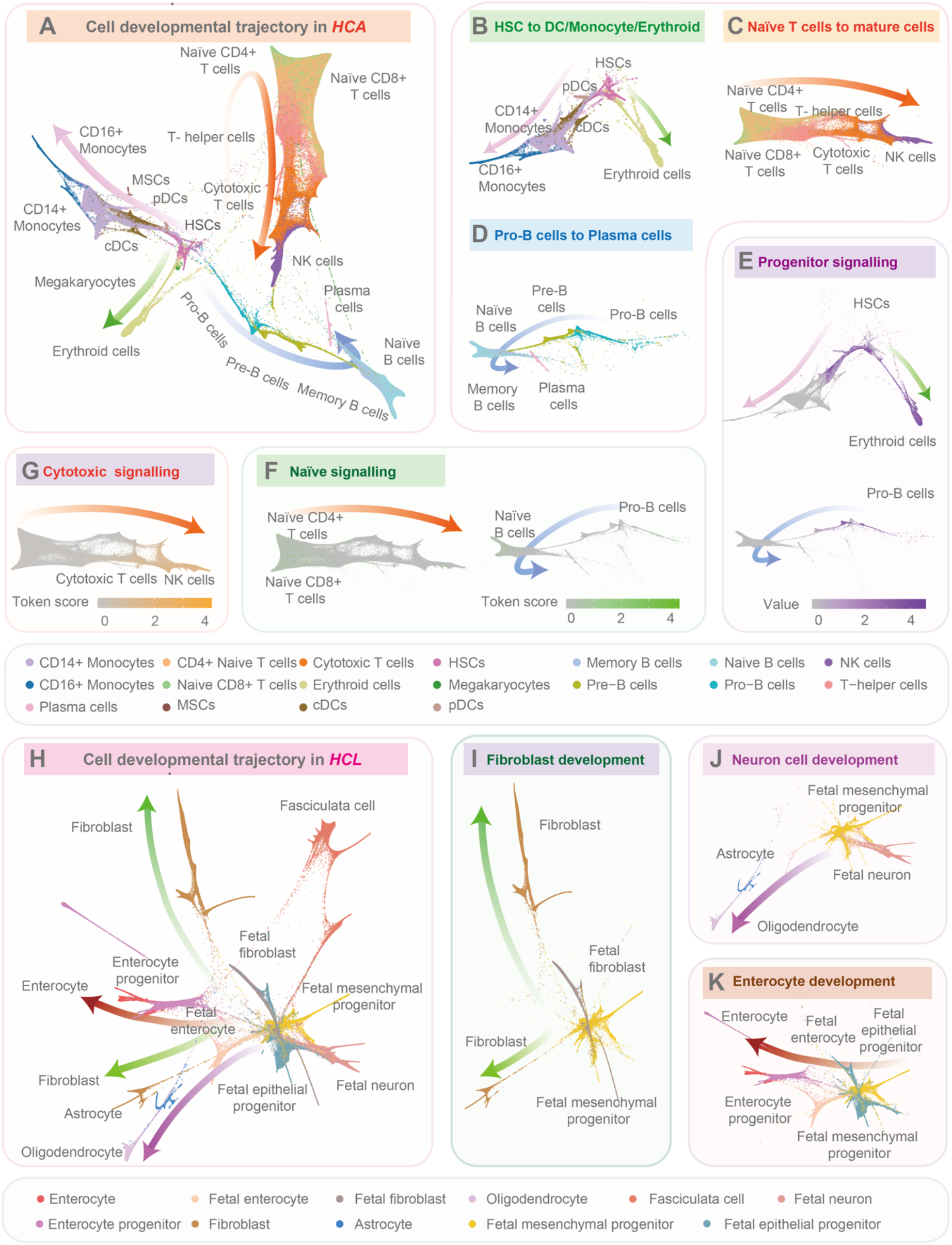
Diffusion pseudo-time analysis on the *HCA* and *HCL* datasets. **(A)** The diffusion map of *HCA* dataset. **(B)** hematopoietic stem cells (HSCs) to erythroid cells or dendritic cells (DCs) and monocytes; **(C)** naïve T cells to cytotoxic T cells and nature killer (NK) cells; **(D)** pro-B cells to plasma cells. **(E-G)** Cell state signatures for progenitor signaling, naïve signaling and cytotoxic signaling. **(H-K)** The diffusion map of *HCL* dataset and its main branches.

On the *HCL* dataset, the developmental tree depicted three differential trajectories of fetal mesenchymal progenitor cells into different mature cell types (**Figure 4H**) with fetal cells at the center of the landscape. The fetal mesenchymal progenitor cells are differentiated into biologically functional fibroblasts (**Figure 4I**), enterocytes (**Figure 4J**), astrocytes and oligodendrocytes (**Figure 4K**).

### Clinical significance of *tGPT* in bulk sequencing sample

Here, we demonstrated that *tGPT* is able to capture clinically significant patterns. On the *TCGA* dataset, we found that the importance scores are varying considerably for different attention heads among different layers. The importance score patterns can cluster different cancer types into distinct groups in that cancer of the same tissue-of-origin are closely related whereas cancers of different origins are well separated (See **Methods**, **Figure 5A**). For example, skin cutaneous melanoma (SKCM) and uveal melanoma (UVM), glioblastoma multiforme (GBM) and brain lower grade glioma (LGG) are respectively located in the same clustering branches. In addition, we examined the association between attention head entropy and molecular alteration events (See **Methods**). There are several attention heads exhibited significant association with tumor mutation burden (TMB) in the TCGA pan-cancer cohort and specifically in bladder urothelial carcinoma (BLCA), LUAD and LUSC (**Figure 5B**). We observed that attention heads also showed significant association with *TP53* mutations at the pan-cancer level and across 9 cancer types (**Figure 5C**). There are also attention heads exhibited significant association with homologous recombination deficiency (HRD) and genome doubling (**Figure 5D** and **E**) at the pan-cancer level. The association of attention heads with HRD and genome doubling are statistically significant across 4 and 14 cancer types, respectively. Meanwhile, the attention heads exhibited prognostic significance at pan-cancer level (**Figure 5E**) and across 7 cancer types (**Supplementary Figure 16**).

**Figure 5.**
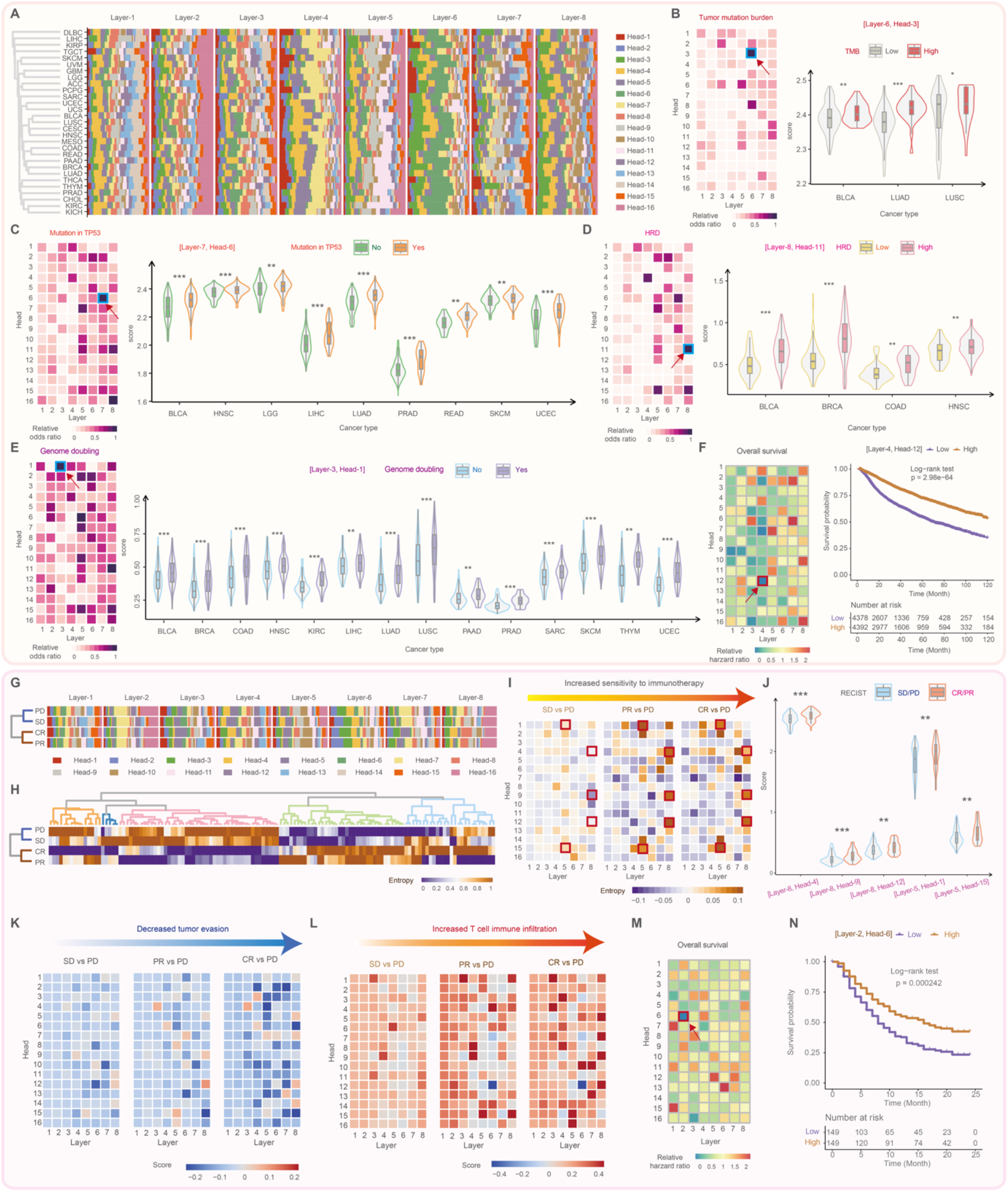
The association of features learned by *tGPT* versus genomic alteration events and clinical phenotype. **(A)** Heatmap representation of attention head importance score across different cancer types on the *TCGA* dataset. Association of attention head entropy versus tumor mutation burden (**B**), *TP53* mutation (**C**), homologous recombination deficiency (**D**), genome doubling **(E)** and overall survival (**F**) on the TCGA cohort. **(G** and **H)** Heatmap representation of attention head importance and entropy on the urothelial carcinoma stratified by RECIST response. CR, complete response; PR, partial response; SD, stable disease; PD, progress disease. **(I)** The varying entropy patterns from SD to PR to CR with PD as baseline. **(J)** Exemplified violin plots depicting attention head entropy in SD/PD versus CR/PR. (**K**) The varying of patterns of tumor evasion signature from SD to PR to CR with PD as baseline. (**L**) The varying of patterns of T cell infiltration signature from SD to PR to CR with PD as baseline. **(M)** Association between attention head entropy and overall survival on the urothelial carcinoma dataset. (**N**) Exemplified survival curves stratified by attention head entropy.

In addition, we examined the attention head patterns in relation to immunotherapy in an immune checkpoint block (ICB) clinical trial of urothelial carcinoma consisted of 298 patients: 25 patients with CR, 43 with PR, 63 with SD and 63 with PD (See **Methods**). We found that importance and entropy scores are distinguishable amongst patients with different therapeutic outcome (**Figure 5G and H**). We observed gradually varying entropy values from SD to PR to CR by taking the PD baseline (**Figure 5I**) and significant difference among 5 attention heads in patients with CR/PR versus SD/PD (**Figure 5J**). We quantified expression signatures such as tumor evasion and T cell immune infiltration attended by different attention heads (See **Methods**). By taking PD as baseline, we observed a gradually decreasing patterns of tumor evasion and increasing patterns of T cell immune infiltration from SD to PR to CR (**Figure 5K** and **L**). The attention heads also exhibited prognostic significance in this clinical trial (**Figure 5M** and **N**).

## Discussion

Efficient integration of increasingly large-scale single-cell transcriptomes is urgently needed. Here, we introduced a conceptually simple approach towards the integration of unlimited number of single-cell transcriptomes and its potential clinical translational relevance. The paradigm underpinning *tGPT* in essence is to predict the occurrence of a given gene with its previous context. We developed *tGPT* on a super large-scale single-cell transcriptomes which consists of 22.3 million cells and systematically evaluated its representation learning ability on different single-cell analysis tasks. We note that *tGPT* was insensitive to batch effect and achieved competitive accuracies as compared with benchmark tools. The purpose of this study is to verify the validity of this new paradigm in deciphering large-scale transcriptome data, especially at the level of single-cell atlas. In addition, we showed that the pretrained *tGPT* model can be applied to bulk tissue sequencing samples to extract a variety of features exhibiting significant association with genomic alteration and potential clinical application.

Artificial intelligence is undergoing a paradigm shift and the pretraining models based on transformer are becoming de facto standard in natural language processing and computer vision, achieving state-of-the-art across a wide range of tasks such as natural language understanding, image classification, video and audio recognition^11^. Representative pretraining models include BERT^14^ and GPT^15^. The advantage of these pretraining models lie in its ability to assimilate real-world information from super large-scale unlabeled and high-dimensional data. This advantage brings an attractive solution for deciphering single-cell transcriptomes as millions of cells have been sequenced, which exemplified by 22.3 million cells collected in our study. This number is expected to increase expotentially in years ahead. There is no analytical tool that are designed and evaluated on such large volume of data. The high expressivity and scalability of transformer enable *tGPT* to learn rich representation from transcriptomes in a self-supervised manner. The high clustering performance in single-cell cluster delineation is probably attributable to better feature representation learned by *tGPT*. In addition, feature representation from *tGPT* is insensitive to batch effect as the acceptance rate of *kBET* derived from *tGPT* is evenly distributed among the other tools that explicitly used batch information for batch-correction. This is probably due to the use of rankings of top expressing genes rather than actual expression levels by *tGPT*. *tGPT* is quite different from the other integration tools^28,29,23^ as the laters use the actual expression levels of highly variable genes (HVGs) and the batch information. The independence of *tGPT* on batch information makes it attractive for super large-scale transcriptome integration since the batch information is not always available and often neglected by researchers.

The clustering performance in delineating single-cell clusters is robust with respect to the number of top expressing genes used and feature representation extracted from different *tGPT* transformer layers. The clustering metrics obtained from 62 top-expressing genes are comparable to the use of 126 top-expressing genes (**Supplementary Figure 1**). This suggested that the rankings of 62 top-expressing genes are sufficient for cell cluster definition. The idea underpinning *tGPT* is to predict the occurrence of a gene in the context of the occurrences of its preceding neighbors. This type of pretraining is not directly related to cell clustering. This does not guarantee that feature representation learned by the last transformer layer could give rise to better clustering as compared with representation learned by all its previous layers. In our evaluation, the cluster metrics obtained from different transformer layers are comparable and consistently better than the embedding layer (**Supplementary Figure 1**). In addition, we observed that cell-type specific genes have high attribution scores albeit only the rankings are used during pretraining. This finding to some extent can explain why features derived from *tGPT* could lead to high performance in cell clustering.

A new finding emerged from our study is that the pretrained *tGPT* model can be applied to bulk tissues. On the *GTEx* dataset, the feature representations of different organs extracted from *tGPT* can divide samples into distinct clusters, aligning with organs. On the *TCGA* dataset, we observed that different cancer types are well separated and cancers of the same origins are more closely related, which is consistent with previous report^39^. Additionally, the feature patterns of TCGA samples exhibited consistent and significant association with genomic alterations. This indicated that rankings of top-expressing genes carry information about alterations in tumor tissues. Meanwhile, the feature patterns derived from *tGPT* are distinctive among patients with different treatment outcomes for immunotherapy. Token together, our finding would facilitate translational research enabled by super large-scale transcriptomes.

We focused two main directions of *tGPT* for future development. Firstly, *tGPT* can be used to generate large-scale reference mapping with the availability of large-scale disease reference datasets and phenotypes. Secondly, *tGPT* can be further investigated for clinical application such as treatment guiding and prognostic prediction.

In summary, we systematically verified a new, simple and effective analytical paradigm for super large-scale transcriptome analysis and its implications in clinical translation.

## ACKNOWLEDGEMENTS

We are grateful for researchers for their generosity to made their data publicly available. This work was supported by the National Natural Science Foundation of China (Grant No. 31801117 to X.L., 31900471 to M.Y. and 82073287 to Q.Z.), National Key Research and Development Program of China (Grant No. 2021YFC2500400 to K.C.), Program for Changjiang Scholars and Innovative Research Team in University in China (Grant No. IRT_14R40 to K.C.) and Tianjin Municipal Health Commission Foundation (Grant No. RC20027 to Y.L).

## AUTHOR CONTRIBUTIONS

Xiangchun Li and Kexin Chen designed and supervised the study; Xiangchun Li and Hongru Shen performed data analysis and wrote the manuscript; Xiangchun Li developed the model; Xiangchun Li, Hongru Shen, Xilin Shen, Chao Zhang, Dan Wu, Mengyao Feng, Jiani Hu, Jilei Liu, Yichen Yang, Yang Li, Meng Yang, Wei Wang and Qiang Zhang collected data; Hongru Shen, Xiangchun Li, Kexin Chen and Jilong Yang revised the manuscript.

## DECLARATION OF INTESTS

The authors declare that they have no conflict of interest.

## Data and code availability

All the gene expression matrices were downloaded from public databases. The source list of these datasets was provided in **Supplementary Table 1**. We will release the pretrained *tGPT* model and its training code at Github soon.

## Methods

### Input preprocessing

The input sequence list of top-expressing genes was obtained via descending sorting. The input to *tGPT* was formulated as [*<s>*, *G_1_*, *G_2_*, *G_3_*, …, *<e>*], where *G_1_*, *G_2_* and *G_3_* are gene symbols and *<s>* and *<e>* are two special tokens respectively added to the start and end of the input sequence. The input sequence is padded with special token *<pad>* if its length is less than a predefined value. The input sequence list is truncated if its length exceeds the predefined value. We evaluated a length of 64 and 128 in this study. The dictionary used by *tGPT* consists of 20706 protein-coding genes.

### The architecture of *tGPT*

***Embedding layer*** transforms the input gene symbols into a real-value matrix that carries the information on gene token embedding and position encoding. The gene token embedding was obtained via an embedding layer (parameterized as *W_e_*) that maps the indices of input genes obtained from the gene symbol dictionary to real-value space. The position encoding (parameterized as *W_p_*) carries information on the sorted gene rankings. For an input sequence *U* = {*G*_−*k*_, … *G*_−1_}, where *k* is the width of context window, the embedding layer injects position encoding onto gene token embedding as:

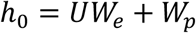

***Transformer decoder blocks*** applies multi-headed masked self-attention over the input embeddings followed by position-wise feed-forward layers, then through a *softmax* layer. *tGPT* use a multi-layer of *transformer decoder^17^*.

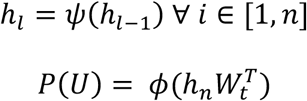

where *ψ* is the transformer decoder block and *ϕ* is the softmax layer, and *W_t_* is the embedding matrix of the *l^th^* decoder block.

***Masked Self-Attention*** is a variant self-attention mechanism^40^. Each attention head adopts the scale dot-product attention to map a query and a set of key-value pairs to an output. The input consists of query and key of dimension *d_k_*, and value of dimensions *d_v_*. Self-attention is calculated as dot products of the query (*Q_i_*) with key (*K_i_*) divided each by 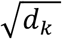 and multiply with value (*V_i_*) after *sofmax* transformed^41^:

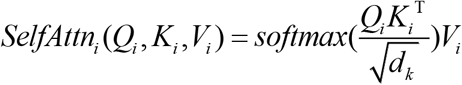

Masked self-attention is implemented with the aid of attention mask. It basically always scores the future tokens as 0 so *tGPT* cannot pick from future. The multi-head self-attention is formulated as:

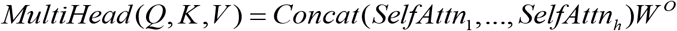

where 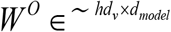 denotes the learned output projection matrix.

***Position-wise Feed Forward neural network*** is a layer with fully-connected feed-forward layer. This layer consists of two linear transformations with a *ReLu* activation function in between:

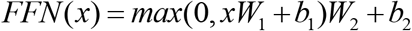

where *W*_1_ and *W*_2_ are weight matrices and *b*_1_ and *b*_2_ are the bias.

### Training scheme

*tGPT* was pretrained with a batch-size of 64 for 100 epochs. We used Adam with *β_1_* = 0.9, *β_2_* = 0.95, weight decay of 0.01 and a learning rate of 0.003. The learning rate is warmed up for four epochs, and then decays to 0 following a cosine schedule^12^. *tGPT* was trained with *PyTorch* (version 1.7.1) and *transformers* (version 4.10.0) on NVIDIA DGX A100 with 8 GPUs each with 40 Gb memory.

### Clustering on feature representation from *tGPT*

We respectively extracted the feature representations from the embedding layer and 8 different transformer layers. The extracted features were used to construct K-Nearest Neighbors (KNN) graphs for subsequent community detection by Leiden algorithm^42^ implemented in *Scanpy* (version 1.8.1). We performed grid-search to identify optimal values of two parameters *n_neighbors* and *resolution* that are the most relevant for clustering. The value of *n_neighbors* examined was ranged from 5 to 100 with step of 5. The value of *resolution* examined was ranged from 0.1 to 2 with step of 0.2. The uniform manifold approximation and projection (UMAP) visualization^43^ is used.

### Features derived from self-attention

***Entropy*** of the self-attention matrices for a given input sequence is calculated as^44^:

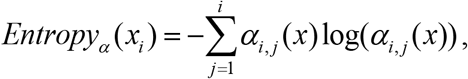

where *α* is the self-attention matrix and *α_i,j_* is the attention weight between the *i^th^* and *j^th^* tokens. We averaged the entropy of all cells in a cluster to derive a cluster-level entropy.

***Head importance score***^45^ is defined as the influence of input on head output. It is calculated via gradient backpropagation, formulated as:

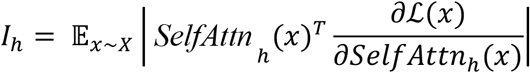

where *x* is the input sequence and 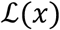 is the corresponding loss given the input. *I_n_* is high score while *SelfAtt_n_*(*x*) is liable to have a large effect on the model.

***Token attribution score*** is defined as the norm of the learned token features (*x_i_*) extracted from *tGPT*, which is defined as:

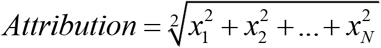

### Attention analysis in relation to signaling

We define an attention-based pathway signaling score in a similar way as^46^:

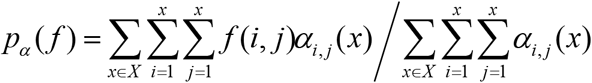

where *α_i,j_* is the attention weight between the *i^th^* and *j^th^* gene. For a given gene signature, we set *f*(*i*, *j*) = *1* if the *i^th^* gene or *j^th^* gene occurs in that gene set.

### Dataset collection

We collected the transcriptomes of 22.3 million single-cells (**Supplementary Table 1**), 9318 bulk tissue transcriptomes of TCGA cohort from the supplemental data of pan-cancer immune landscape study^22^, 11,688 bulk tissue transcriptomes from *GTEx* database^21^ and 298 bulk tissue transcriptomes from the clinical trial study on immunotherapy for urothelial carcinoma^47^. We discarded mitochondrial genes, ribosomal genes and non-protein coding genes for the single-cell data. Four single-cell and two bulk tissue sequencing datasets are used in downstream evaluation of *tGPT*:

#### Human Cell Atlas Census of Immune Cells (HCA)

Bone marrow cells (n = 282,588) from 64 healthy donors in *Human Cell Atlas* (*HCA*) project. The data are subjected to 10x sequencing protocol^18^ and contained 18 cell types such as hematopoietic stem cells (HSCs), mesenchymal stem cells (MSCs), erythrocytes, megakaryocytes and different kinds of immune cells.

#### Human cell Landscape (HCL)

HCL dataset includes 586,135 human cells obtained from a Chinese Han population^19^, the dataset encompasses samples of fetal and adult tissue and covered 60 human tissue types, and are subjected to Microwell-seq protocol.

#### Tabula Mursi

The *Tabula Muris* dataset (n = 54,865) is consisted of single-cells sorted by FACS from Mouse Cell Atlas^4^ across 20 different organs subjected to 10x and Smart-seq2 sequencing protocols.

#### The Cancer Genome Atlas (TCGA)

The *TCGA* dataset is consisted of 9,318 bulk samples with primary cancer and matched normal samples spanning 33 cancer types.

#### Genotype-Tissue Expression Project (GTEx)

The *GTEx* dataset includes 11,688 bulk samples across 30 organs obtained from healthy donors.

***Known marker genes*** of different cell types are curated from CellMarker database^48^: plasma cell (*MZB1*), DCs and monocytes (*CST3*, *FCER1A*, *IRF7*, *CD14* and *FCGR3A*), megakaryocyte (*PPBP* and *PF4*), B cell (*CD79A*, *CD79B* and *MS4A1*) NK and cytotoxic T cell (*NKG7*, *FGFBP2*, *GNLY*, *GZMA*, *GZMB* and *PRF1*).

***Cell state signatures*** are curated from CellMarker database^48^, including progenitor signaling (*STMN1*, *TUBA1B* and *HIST1H4C*), naïve signaling (*CCR7*, *LEF1* and *SELL*) and cytotoxic signaling (*GZMA*, *CD8A*, *CD8B*, *GZMB*, *PRF1*, *IL2*, *GNLY*, *GAMK*, *IFNG* and *NKG7*).

***T cell infiltration signature*** is obtained from CellMarker database^48^; it consists of *CD3D*, *CD3E* and *CD8A*.

***Tumor evasion signature*** is curated from the Figure 1 of a previous study^49^.

### Diffusion pseudo-time maps construction

We constructed the diffusion pseudo-time maps using package *Pegasus*^24^ (v1.4.3), and the cell trajectory was visualized with force-directed layout embedding (FLE) algorithm^50^. We set *δ* and *nδ* to its default values: *δ* = 2.0 and *nδ* =5,000.

Firstly, we used the features obtained from last transformer decoder blocks to construct affinity matrix of cells *W_n×n_*, and the top-*k* nearest neighbor cells were find by community detection algorithm^51^ and the HNSW algorithm^52^, and the formula of affinity matrix is define as:

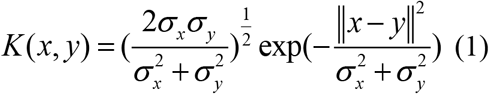

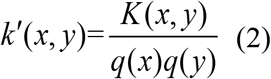

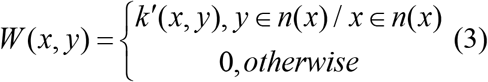

The formula (1) represented the distance between cell-*x* and cell-*y*, σ_x_ is the *x*’s local kernel width, *x* and *y* are features of last transformer decoder block for cell-*x* and cell-*y*. The affinity matrix *W* was calculated as the density-normalized kernel according to formula (3).

We then calculated the Markov chain transition matrix *P* and the symmetric transition matrix *Q* as the formula:

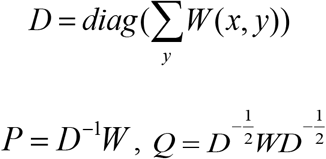

The symmetrical matrix *Q* can be decomposed as *UAU^T^*. Let 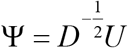 A family with parameter timescale of *t* for approximated diffusion maps 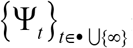 is defined as:

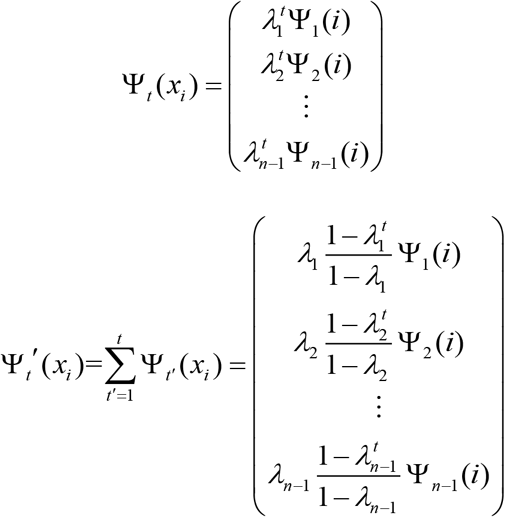

### Benchmark methods

We also performed single-cell analysis using *Scanpy* (version 1.6.0), *Pegasus* (version 1.4.3) and *scVI* (version 0.13.0). Batch-correction was performed with *MNN* (version 1.8.0)^26^, *Combat* (version 1.8.0)^53^, *Harmony* (version 0.1.6)^27^, *Seurat* (version 3.1.5)^28,29^, *Pegasus* (version 1.4.3)^24^, *Scanorama* (version 1.7.1)^31^, *DESC* (version 2.1.1)^34^, *iMAP* (version 1.0.0)^33^, *scVI* (version 0.13.0)^8^, *scArches* (version 1.7.0)^32^, *BBKNN* (version 1.7.1)^30^.

***Scanpy*** is a comprehensive toolkit for analyzing single-cell transcriptome. We first filtered out cells with the number of expressing genes < 200 or *mitochondrial counts* > 30%. We used the function *scanpy.pp.highly_variable_genes* to selected highly variable genes by setting *max_mean* to 3 and *min_mean* to 0.0125, which are the default values. We then applied clustering pipeline and grid-search to perform single-cell clustering on KNN graph. The UMAP is used for visualizing clustering result.

***scVI*** is a deep generative model for mining the single-cell omics data. We filtered out cells with the number of expressing genes < 200 or *mitochondrial counts* > 30%, and selected HVGs with *scanpy.pp.highly_variable_genes* by setting *max_mean* to 3 and *min_mean* to 0.0125. We used the default parameter of *scVI* to extract the 10 latent features. These latent features were used to construct KNN graphs for community detection by Leiden algorithm^42^.

***Pegasus*** is complete single-cell analysis pipline that is efficient on large datasets. We used the recommended parameters: *min_genes* of 500, *max_genes* of 6000, and *percent_mito* of 10. We identified the robust genes with the default *percent_cells* of 0.05. Single-cell clustering was performed on KNN graph followed by Leiden algorithm^42^ for community detection.

### Clustering and batch-effect metrics

We used Adjusted Rand Index (*ARI*), Normalized Mutual information (*NMI*) and Fowlkes-Mallows Index (*FMI*) to measure clustering performance. We used the *kBET* acceptance rate^35^ as a measurement of batch-effect. The clustering metrics of *ARI*, *NMI* and *FMI* were calculated with sklearn (version 0.21.2). *kBET* acceptance rate is computed with *Pegasus* (version 1.4.3).

***ARI*** is calculated based on the contingency table summarizing the truth labels and clustering, and the rows and columns represent truth and clustering labels in the contingency table, respectively. The formula is as follows:

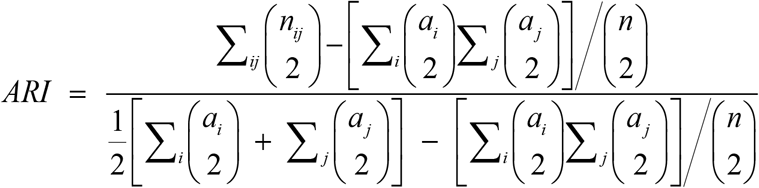

where *n_ij_* denoted the numbers of cell in common between clustering labels and truth labels, *a_i_* the sum of *i^th^* row and *a_j_* the sum of *j^th^* column of the contingency table.

***NMI*** is also used to measure the similarity between the clustering labels and actual labels. We assumed that the clustering labels and actual labels of *N* cells are *U* and *V*, and the entropy of *U* and *V* is as the following formula:

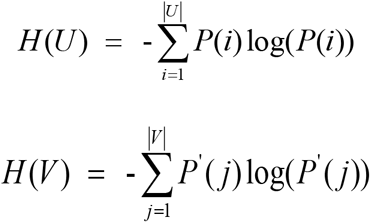

where *p*(*i*) = |*U_i_*| / *N* is the probability that a cell picked at random from *U* falls into *U_i_*, *p′*(*j*) = |*V_i_*| / *N* is the probability that a cell picked at random from *V* falls into *_V_j*. We then calculated the mutual information (MI) between *U* and *V*, and normalized the mutual information:

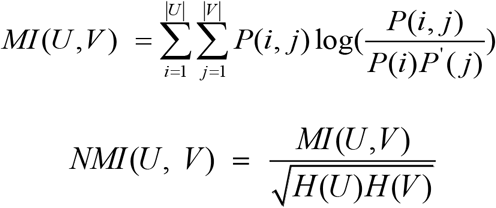

where *p*(*i*, *j*) = |*U_i_* ∩ *V_j_*| / *N* is the probability that a cell picked at random falls into classes *U_i_* and *V_j_*.

***Fowlkes-Mallows Index (FMI)*** is used to measure the consistency between clustering results and real category, and the range of index is from 0 to 1. The *FMI* metric is denfined as the geometric mean between of the precision and recall:

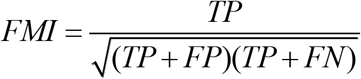

where *TP* is true positive, FP false positive, FN false negiative.

***kBET acceptance rate*** is a measurement of batch effect. We assumed that the dataset of cells with batches of *m*, and there are *n_j_* cells in batch *j*. The batch mixing frequency denotes as:

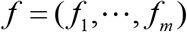

where 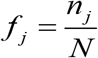. Then, we calculated the number of neighbors of cell-*i* belonging to batch *j* is 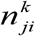. Its χ^2^ test statistic and *p*-value with degrees of (m-1) are defined as follows:

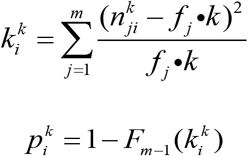

where *F*_*m*−1_ (*x*) represents the cumulated density function. The *kBET* acceptance rate is defined as the percentage of cells that accept the null hypothesis at significance level α as follows:

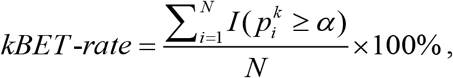

*I(x)* is the indicator function where *I(x)* = 1 if *x* > 0 otherwise *I(x)* = 0. We used *Pegasus* (v1.4.3) to calculate the *kBET* acceptance rate by setting *K* and *α* to 5 and 0.01, respectively.

---

**Supplementary Figure 1. Radar charts illustrating the clustering performance achieved by feature representations extracted from different layers of *tGPT* for the top 62 (A) and 126 (B) expressing genes on *HCA*, *HCL, Tabula Muris, Macaque Retina, GTEx, and TCGA* datasets**.

**Supplementary Figure 2. The clustering performance with grid search for resolution and number of neighbors for the top 62 (A) and 126 (B) expressing genes among feature representations extracted from different layers on the *HCA* dataset**. Contour maps depict different cluster metrics (i.e. *NMI*, *ARI* and *FMI*) with respect to different values of *Resolution* and *N-neighbors*.

**Supplementary Figure 3. The clustering performance with grid search for resolution and number of neighbors for the top 62 (A) and 126 (B) expressing genes among feature representations extracted from different layers on the *HCL* dataset**. Contour maps depict different cluster metrics (i.e. *NMI*, *ARI* and *FMI*) with respect to different values of *Resolution and N-neighbors*.

**Supplementary Figure 4. The clustering performance with grid search for resolution and number of neighbors for the top 62 (A) and 126 (B) expressing genes among feature representations extracted from different layers on the *Tabula Muris* dataset**. Contour maps depict different cluster metrics (i.e. *NMI*, *ARI* and *FMI*) with respect to different values of *Resolution* and *N-neighbors*.

**Supplementary Figure 5. The clustering performance with grid search for resolution and number of neighbors for the top 62 (A) and 126 (B) expressing genes among feature representations extracted from different layers on the *Macaque Retina* dataset**. Contour maps depict different cluster metrics (i.e. *NMI*, *ARI* and *FMI*) with respect to different values of *Resolution* and *N-neighbors*.

**Supplementary Figure 6. The clustering performance with grid search for resolution and number of neighbors for the top 62 (A) and 126 (B) expressing genes among feature representations extracted from different layers on the *GTEx* dataset**. Contour maps depict different cluster metrics (i.e. *NMI*, *ARI* and *FMI*) with respect to different values of *Resolution* and *N-neighbors*.

**Supplementary Figure 7. The clustering performance with grid search for resolution and number of neighbors for the top 62 (A) and 126 (B) expressing genes among feature representations extracted from different layers on the *TCGA* dataset**. Contour maps depict different cluster metrics (i.e. *NMI*, *ARI* and *FMI*) with respect to different values of *Resolution and N-neighbors*.

**Supplementary Figure 8. UMAP visualization on different datasets obtained from different methods**.

**Supplementary Figure 9. The full annotation of UMAP visualization of different methods on the *HCL* dataset**. The *NMI* metric and annotation of cells are shown.

**Supplementary Figure 10. The full annotation of UMAP visualization of different methods on the *Tabula Muris* dataset**. The *NMI* metric and annotation of cells are shown.

**Supplementary Figure 11. Radar charts illustrating the best clustering metrics for different methods across different datasets obtained from grid search**. *ARI*, Adjusted Rand Index; *NMI*, Normalized Mutual information; *FMI*, Fowlkes-Mallows Index.

**Supplementary Figure 12. The clustering performance with grid search for resolution and number of neighbors for *Scanpy (A), Pegasus (B),* and *scVI (C)* on the *HCA*, *HCL*, *Tabula Muris* and *Macaque Retina* dataset**. Contour maps depict different cluster metrics (i.e. *NMI*, *ARI* and *FMI*) with respect to different values of *Resolution* and *N-neighbors*.

**Supplementary Figure 13. The UMAP visualization plots of different batch-correction methods on the HCA dataset (A to K) and *kBET* acceptance rate (L)**.

**Supplementary Figure 14. Radar charts illustrating the best clustering metrics for different batch-correction methods obtained from grid search on the *HCA* dataset**. *ARI*, Adjusted Rand Index; *NMI*, Normalized Mutual information; *FMI*, Fowlkes-Mallows Index.

**Supplementary Figure 15. The clustering performance with grid search for resolution and number of neighbors for different batch-correction methods on the *HCA* dataset**. Contour maps depict different cluster metrics (i.e. *NMI*, *ARI* and *FMI*) with respect to different values of *Resolution* and *N-neighbors*.

**Supplementary Figure 16. The survival curves stratified by attention head entropy across multiple cancer types on TCGA dataset**. ACC, Adrenocortical carcinoma; CHOL, Cholangiocarcinoma; KIRC, Kidney renal clear cell carcinoma; READ, Rectum adenocarcinoma; KIRP, Kidney renal papillary cell carcinoma; LGG, Brain Lower Grade Glioma; MESO, Mesothelioma.

**Supplementary Table 1**. Source of data list.

## Notes

### Competing Interest Statement

The authors have declared no competing interest.

